# A Multi-Parameter Analysis of Cellular Coordination of Major Transcriptome Regulation Mechanisms

**DOI:** 10.1101/104901

**Authors:** Wen Jiang, Zhanyong Guo, Nuno Lages, W. Jim Zheng, Denis Feliers, Fangyuan Zhang, Degeng Wang

## Abstract

To understand cellular coordination of multiple transcriptome regulation mechanisms, we simultaneously measured transcription rate (TR), mRNA abundance (RA) and translation activity (TA). This revealed multiple quantitative insights. First, the genomic profiles of the three parameters are systematically different in key statistical features. Sequentially more genes exhibit extreme low or high expression values from TR to RA, then to TA. That is, because of cellular coordination of these regulatory mechanisms, sequentially higher levels of gene expression selectivity are achieved as genetic information flow from the genome to the proteome. Second, the contribution of the stabilization-by-translation regulatory mechanism to the cellular coordination process was assessed. The data enabled an estimation of mRNA stability, revealing a moderate but significant positive correlation between the estimated mRNA stability and translation activity. Third, the proportion of a mRNA occupied by untranslated regions (UTR) exhibits a negative relationship with the level of this correlation, and is thus a major determinant of the mode of regulation of the mRNA. High-UTR-proportion mRNAs tend to defy the stabilization-by-translation regulatory mechanism, staying out of the polysome but remaining stable; mRNAs with little UTRs largely follow this regulation. In summary, we quantitatively delineated the relationship among multiple transcriptome regulation parameters, i.e., cellular coordination of corresponding regulatory mechanisms.

## Background

The genomic sequences are readily available for a large and ever-increasing number of species. These sequences, like English literature, represent static strings of symbols/alphabets (A, T, C, and G). Hence, genomic sequences are often termed as the “book” of life. To some degree, the cell can be considered as the “reader” of the genomic “book”, and the multi-stepped gene expression process as the “reading” process [1-4]. Through the gene expression process, the seemingly simplistic genomic alphabetical strings are selectively and dynamically transcribed into transcriptome sequences, which are in turn translated into amino acid sequences in the proteome – the main machinery that controls biochemical reactions and processes in support of cellular functions. This process is integral to essentially all cellular activity and entails multiple regulatory mechanisms. A complex picture of the relationship among these regulatory mechanisms has recently emerged [5-11].

In some omics experiments, mRNA and protein abundance are measured simultaneously. One lesson we learned is that correlation between the two is not always satisfactory enough for mRNA abundance to be a reliable predictor of protein abundance. This discrepancy had been observed prior to the genomic era [12, 13]. It was confirmed in the yeast *S. cerevisiae* by one of the first simultaneous transcriptome and proteomic measurement [14], and then observed in many other high-throughput studies [5, 9, 10, 15-21].

Transcriptome analysis techniques have also been coupled to conventional experimental protocols to measure other gene expression parameters. Initially micro-array [22-25], and then NGS [26, 27], were coupled to the nuclear run-on technique for genome-wide transcription rate measurement. Additionally, NGS was coupled to metabolic labeling of nascent transcripts to measure transcription rate [10, 28-31]; it is also coupled to RNA polymerase II chromatin immunoprecipitation (ChIP) for the same purpose. These strategies enabled simultaneous transcription rate and mRNA abundance measurement. Once again, some levels of discrepancy were observed in that mRNA abundance was not always a good predictor of transcription rate.

These observed discrepancies among gene expression parameters were a reflection of the complexity of the gene expression process [32], and should be informative for us to unravel the complexity. At the same time nascent RNA and protein are produced, existing RNA and protein are being selectively degraded. The abundance of protein and mRNA represent the balance of the respective production and degradation. Discrepancy among gene expression parameters is considered evidence for some levels of decoupling among transcription, translation, mRNA degradation and protein degradation; that is, the gene expression parameters can be divergently regulated. Given the technical feasibility, multi-parameter approaches are being used to study the discrepancy and glean out fundamental gene expression regulation principles. Such studies will potentially lead to more efficient gene expression analysis strategies that generate more informative data.

Such multi-parameter approaches should be especially fruitful for transcriptome analysis. The polysome profiling analysis utilizes NGS to quantify polysome-associated mRNAs, i.e., actively translating mRNAs, thus enabling genome-wide analysis of translation activity. Similarly, the ribosome profiling analysis utilize NGS to quantify mRNA fragments protected from RNase digestion by the ribosome [33, 34]. Thus, all techniques are in place for genome-wide integration of transcription rate (*i.e.*, GRO-seq), mRNA abundance (RNA-seq) and mRNA translation activity (*i.e.*, polysome profiling). This will generate an integrative view of the transcriptome and its dynamic regulation, i.e., how the multiple transcriptome regulatory mechanisms are coordinated.

Additionally, such data is needed as a platform to study mediation of post-transcriptional regulation by mRNA untranslated regions (UTR), where regulatory signals for post-transcriptional regulation are embedded. It is well documented that mRNA UTRs are responsible for mRNA stability and translation control. They contain binding sites for microRNA and many regulatory RNA-binding proteins. They are common in mammalian mRNAs. Human mRNAs, on average, have ∼1000 nucleotide long UTRs (∼800 nucleotide 3’- and ∼200 nucleotide 5’-UTRs). Systematic functional study of the UTRs, however, awaits multi-parameter datasets that enables simultaneous study of mRNA stability and translation activity.

Thus, we generated a multi-parameter snapshot of the transcriptome by simultaneous genome-wide measurement of transcription rate (TR), mRNA abundance (RA) and translation activity (TA); we also estimated mRNA stability/degradation by the RA to TR ratio. The data enabled a quantitative delineation of cellular coordination of these regulatory mechanisms. Briefly, the data revealed functional consequences of the cellular coordination activity. We assessed, for the first time, the contribution of the mRNA-stabilization-by-translation regulatory mechanism to transcriptome regulation, as it is known that actively translating mRNA is protected from degradation [35, 36]. Analysis of the data in conjunction with mRNA UTRs revealed further insights into, and the roles of UTRs in, cellular coordination of these transcriptome regulatory mechanisms.

## Results

### Simultaneous measurement of TR, RA and TA

Previously, we have analyzed publicly available genomic datasets, in which multiple gene expression parameters are simultaneously measured [37, 38]. In those studies, we attempted to explain the discrepancy among gene expression parameters, which then seemed mysterious to most scientists, from the perspectives of biochemical pathway/network control and cellular operations. Though the genome-wide measurement techniques have since greatly advanced and many datasets have recently been published [39], we have not seen a dataset that integrate TR, RA, TA and mRNA stability; the translational data in such studies are mass-spectrometry-based, and thus the coverage is not nearly genome-wide. Thus, in the present work, we took advantage of the genome-wide analysis power of NGS and its versatility through successful coupling to a variety of conventional experimental protocols. Our goal is to simultaneously measure TR, RA and TA, that is, to obtain a genome-wide multi-parameter snapshot of the transcriptome, in the HCT116 human cells. The experimental strategy is illustrated in Figure 1. The experiments were done with cells in exponential growth (log) phase (see Materials and Methods for details). We measured RA with the standard RNA-seq method. Simultaneously, we measured genome-wide TR and TA, using the GRO-seq technique and the polysome profiling technique, respectively. We chose GRO-seq for two reasons; first, its higher sensitivity than the RNA Polymerase II ChIP-seq (Pol-II ChIP-seq) analysis; and, second, to avoid the need for label-time calibration associated with metabolic labeling based methods (*i.e.*, 4sU-seq). Nevertheless, it has been shown that 4sU-seq data agree well with Pol-II ChIP-seq, and thus GRO-seq, data [29]. Our selection of TR measurement method should, thus, not matter. The NGS reads were aligned to the human genome with TopHat [40] and the read counts for expressed genes were calculated with the HTSeq-count software [41]. The read counts were then converted into <u>R</u>eads <u>P</u>er <u>K</u>ilo-base Per <u>M</u>illion Mapped Reads (RPKM) values. With a cut-off of 1 RPKM for at least one of the three parameters, 12921 genes were found expressed in the HCT116 cells. As expected, the experimental results are highly repeatable. For all three parameters, respective biological replicates are extremely consisten with each other, with linear regression R-squred values of at least 0.94 (see Methods for detail). The dataset provides a unique opportunity to generate mechanistic insight into cellular cordination of major transcriptome regulatory mechanisms.

**Figure 1:**
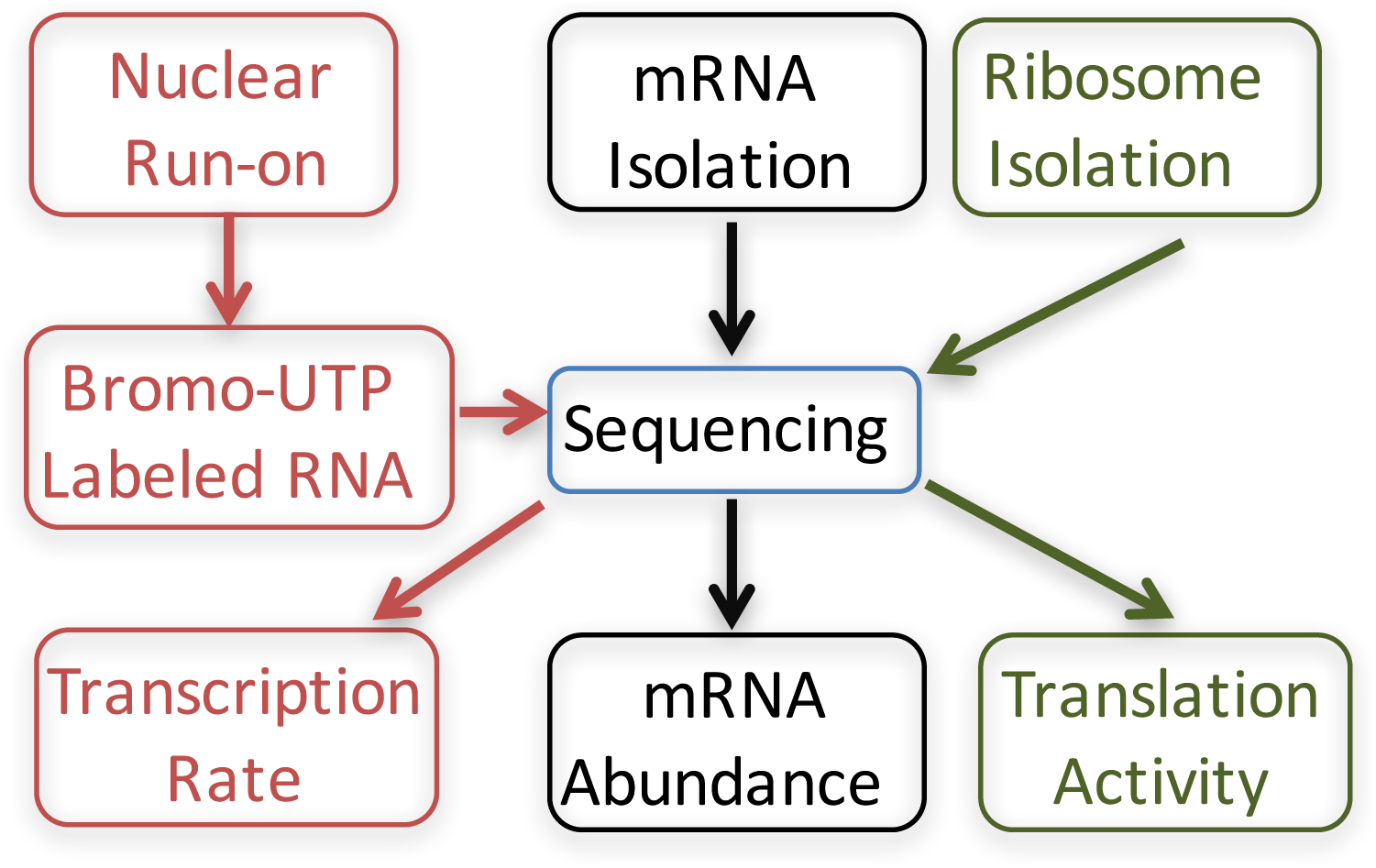
Experimental strategy. Log-phase HCT116 cells were split up into three parts. One part was used to extract total mRNA for RNA-seq analysis to measure steady-state mRNA abundance (RA) (black texts and arrows). One part was used to perform nuclear run-on to generate bromo-UTP labeled nascent RNA for sequencing, that is, GRO-seq analysis to measure transcription rate (TR) (green texts and arrows). The last part was used to isolate and quantify polysome associated mRNA to measure translation activity (TA) (green texts and arrows).

### Comparative analysis of the three gene expression parameters, revealing sequentially higher levels of selectivity from transcription to translation

Comparative analysis of the three gene expression parameters revealed extensive difference among them. Individual pairwise comparison resulted in, as expected, a general trend of good correlation; that is, association of a high value of one parameter with high values of other parameters. However, regression analysis revealed quite dramatic differences, which are way beyond intrinsic experimental noises, among the three parameters (Fig. 2). In Figure 2A, TR and RA are compared in contrast to, in the same scatter plot, a comparison of one RA biological replicate (RA2) against another RA biological replicate (RA1); the two RA replicates illustrate the level of the intrinsic experimental noises. The two RA replicates agree with each other well, with a linear regression slope of about 1 and a low level of dispersion along the regression line. However, the TR-RA regression is dramatically different. The slope of the regression line is only 0.5, suggesting systematic difference between the two parameters. Additionally, as shown in Figure 2B, the TA-RA regression line is also different from the RA2-RA1 line. The change in the slope of the regression line, an increase to 1.11, is not as dramatic. But, statistically, it is highly significant, with a p-value of less than 1E-200 – essentially zero (see Materials and Methods for detail). Thus, systematic discrepancies exist among the three key transcriptome parameters.

**Figure 2:**
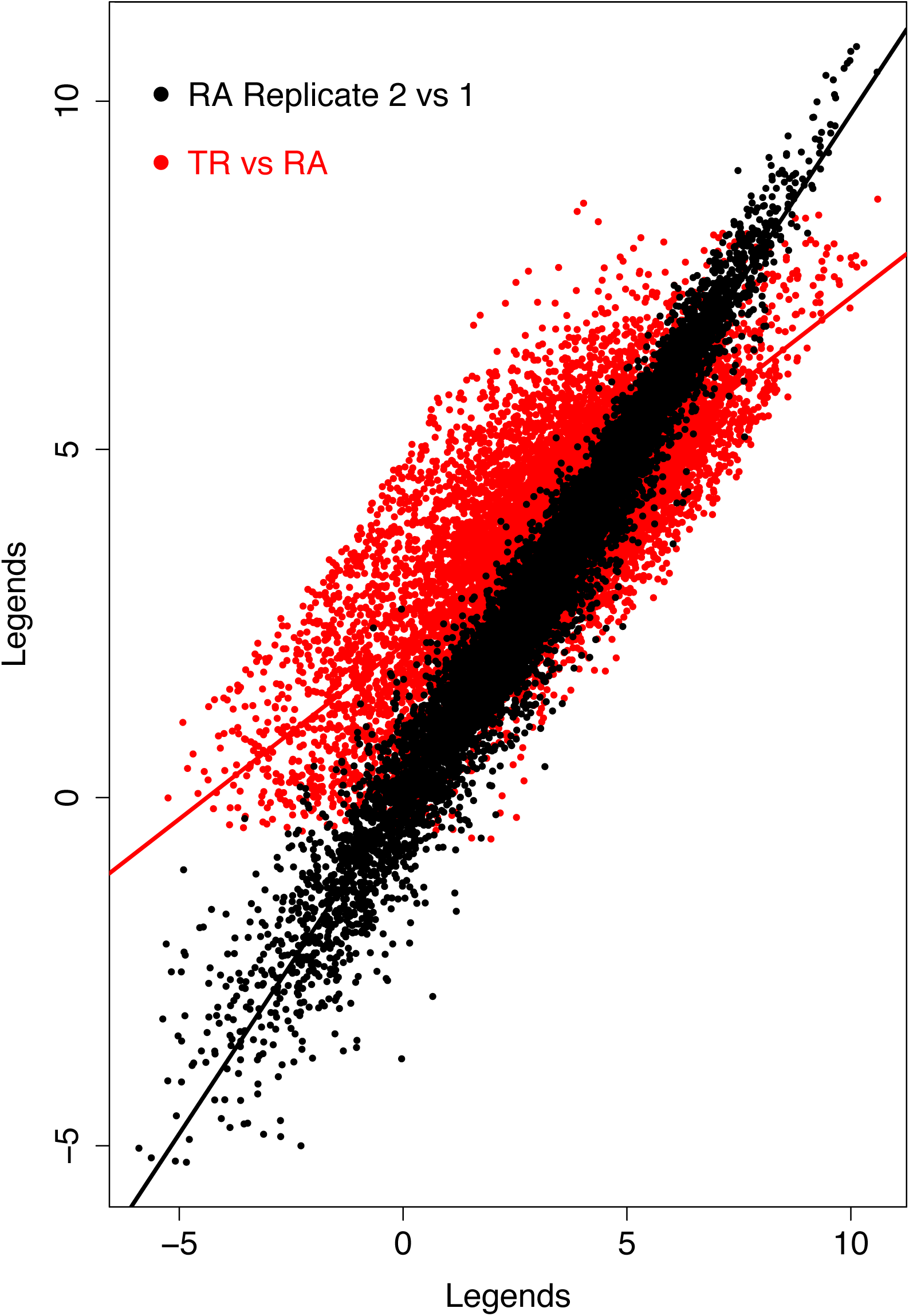
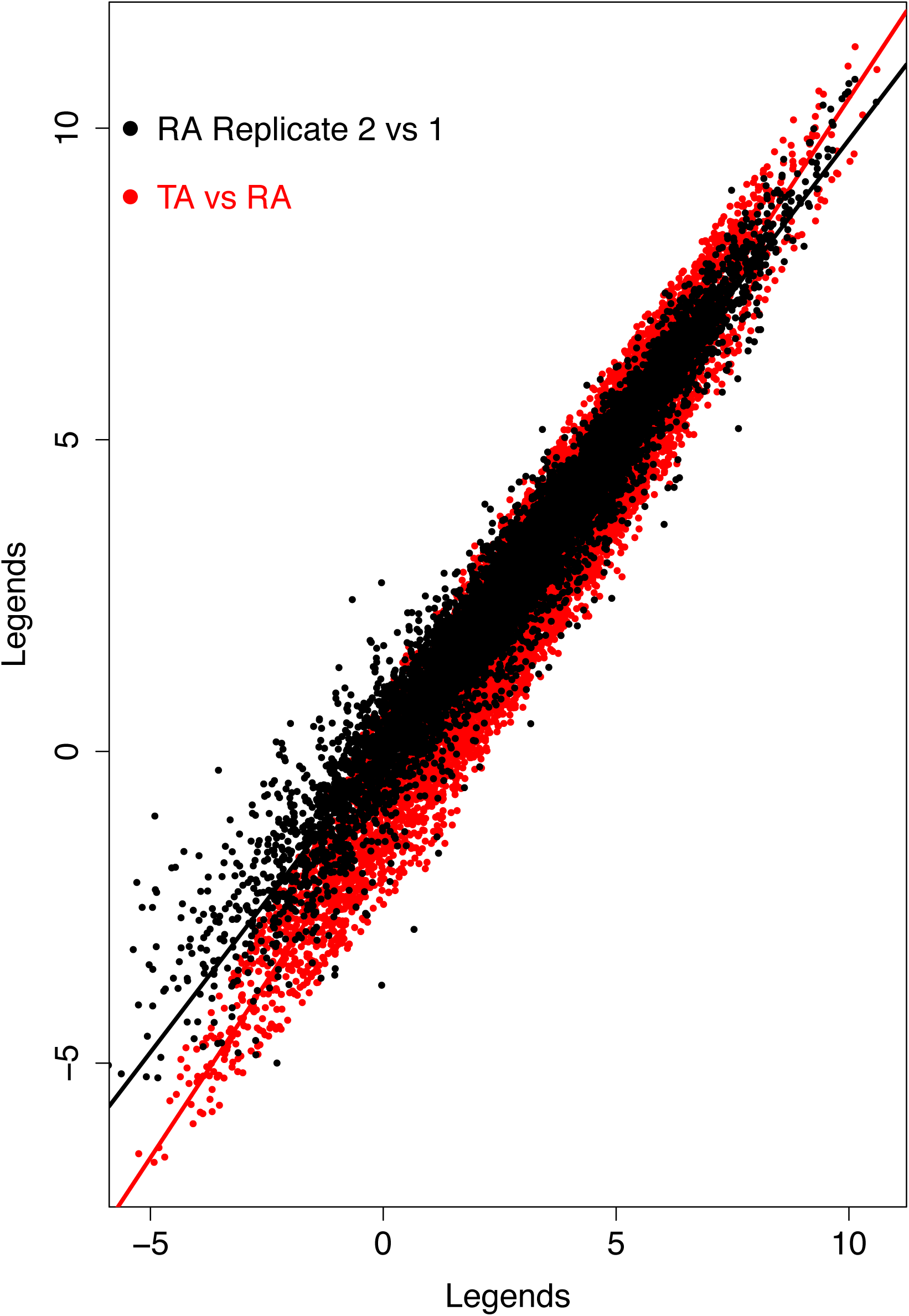
Comparison of TR, RA and TA to illustrate the discrepancy among the three parameters. A. Scatter plot of TR versus RA (red) and a RA experimental replicate versus another RA experimental replicate (black). B. Scatter plot of TA versus RA (red) and a RA experimental replicate versus another RA experimental replicate (black). The same two RA experimental replicates are used in A and B. The linear regression lines are also shown.

We also directly compared the statistical features of the genomic profiles of the three parameters. A systematic trend was observed. The levels of dispersion of the three distributions increase from transcription rate to mRNA abundance, and then to translation activity. Schematically, the trend is also shown in figure 3; quantitatively, this trend is illustrated by the sequential increase of two standard statistical parameters – the standard deviation and the value range – of the three genomic profiles (Table S1). This trend is consistent with the observations in figure 2; the two regression lines in figure 2A indicate that RA tends to have higher values than TR for high expression level genes and lower values than TR for low expression level genes, and thus has higher dispersion than TR; similarly, the two regression lines in figure 2B indicate TA has higher dispersion than RA.

**Figure 3:**
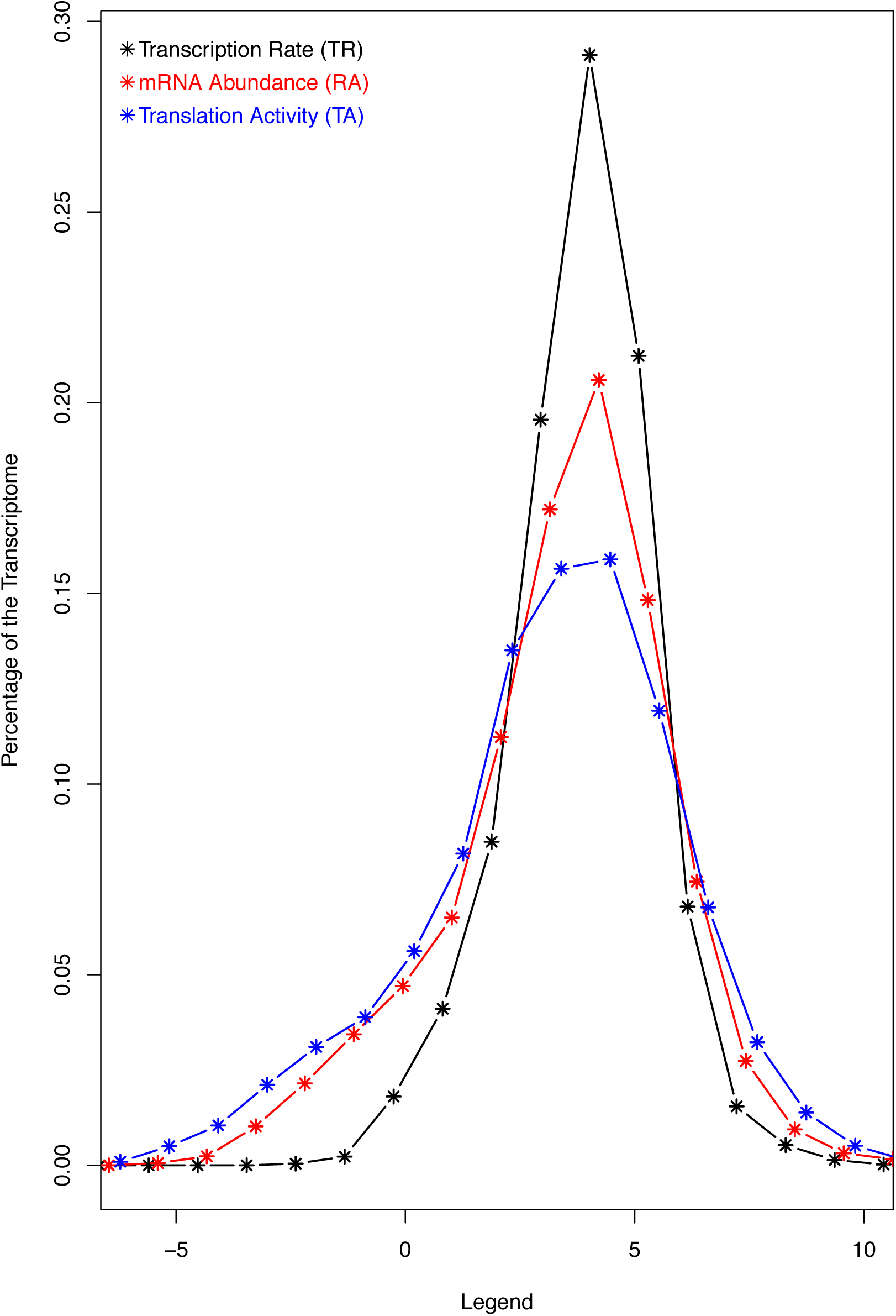
Comparison of the genomic profiles of the three transcriptome parameters – TR, RA and RA. Histograms of TR (black), RA (red) and TA (blue) are shown. The RA and TA histograms are shifted a little bit so that the three histograms have theirs peaks in the same x-axis range, in order to better display the increased levels of dispersion from TR and RA and then to TA.

Thus, both regression analysis and statistical parameters of the distributions demonstrate that more and more genes display extreme (either low or high) parameter values from TR to RA, and then to TA. That is, the functional consequence of cellular coordination of the major transcriptome regulatory mechanisms is sequentially higher level of gene expression selectivity as the genetic information flow in the direction dictated by the Central Dogma. Next, we further dissected our dataset to decipher the underlying mechanisms that give rise to the observed discrepancies and lead to the sequential enhancement of gene expression selectivity.

### Contribution of mRNA-stabilization-by-translation to the sequential enhancement of gene expression selectivity

We tested whether, and to what extent, the TA-RA and the RA-TR discrepancy are related with each other. The TA-RA discrepancy is a reflection of mRNA translation regulation; enrichment of mRNA species with high translation activity in polysome complexes leads to higher TA values than RA values and *vice versa*. In case of TR and RA, the discrepancy results mostly from regulation of both mRNA degradation and, to a much lesser extent, from RNA processing. RNA processing is extensively coupled to transcription [42], and transcription was shown to be on average more than three folds slower than RNA processing [6]. Transcription should be, in most cases, the rate limiting step in mRNA production. Thus, TR, as has been reported, closely correlates with mRNA production rate [6]. By extension, TR-RA discrepancy should be, to large degree, accounted for by mRNA stability control; unstable mRNA (or high degradation rate) leads to lower RA values than mRNA production rate and, in turn, TR values and *vice versa*. Furthermore, it is known that active translation shields mRNAs from degradation, thus stabilizing the mRNA molecules and contributing to the discrepancy between mRNA production rate and RA [35]. In other words, translation activity should be a major determinant of how RA deviates from mRNA production rate and, in light of close correlation between TR and mRNA production rate, the TR-RA discrepancy. Our data provide a unique opportunity, to our knowledge for the first time, for a genome-wide and quantitative assessment of the contribution of this stabilization-by-translation regulatory mechanism to transcriptome regulation. For this purpose, we used the log_2_(TA/RA) and log_2_(RA/TR) log ratios as translation index and TR-RA discrepancy index, respectively. The former is the log ratio between actively translated mRNA abundance and total mRNA abundance, thus a measurement of mRNA translation activity normalized against RA; the latter is the log ratio between total mRNA abundance and transcription rate, and can be operationally considered as a close estimate of mRNA stability.

We hypothesized that the stabilization-by-translation regulatory mechanism should exert a significant effect and lead to positive correlation between the two indices. If our hypothesis is wrong, the two indices should be negatively correlated, since RA is the numerator in the TR-RA index and denominator in the translation index. However, our experimental results turned out to be the contrary and, thus, support our hypothesis. As shown in Figure 4A, a dot-plot of the two indices illustrates an overall positive relationship, with a correlation coefficient of 0.39; the linear regression line is also shown to quantify the relationship, with a slope of 0.19. In other words, an overall positive correlation was observed.

**Figure 4:**
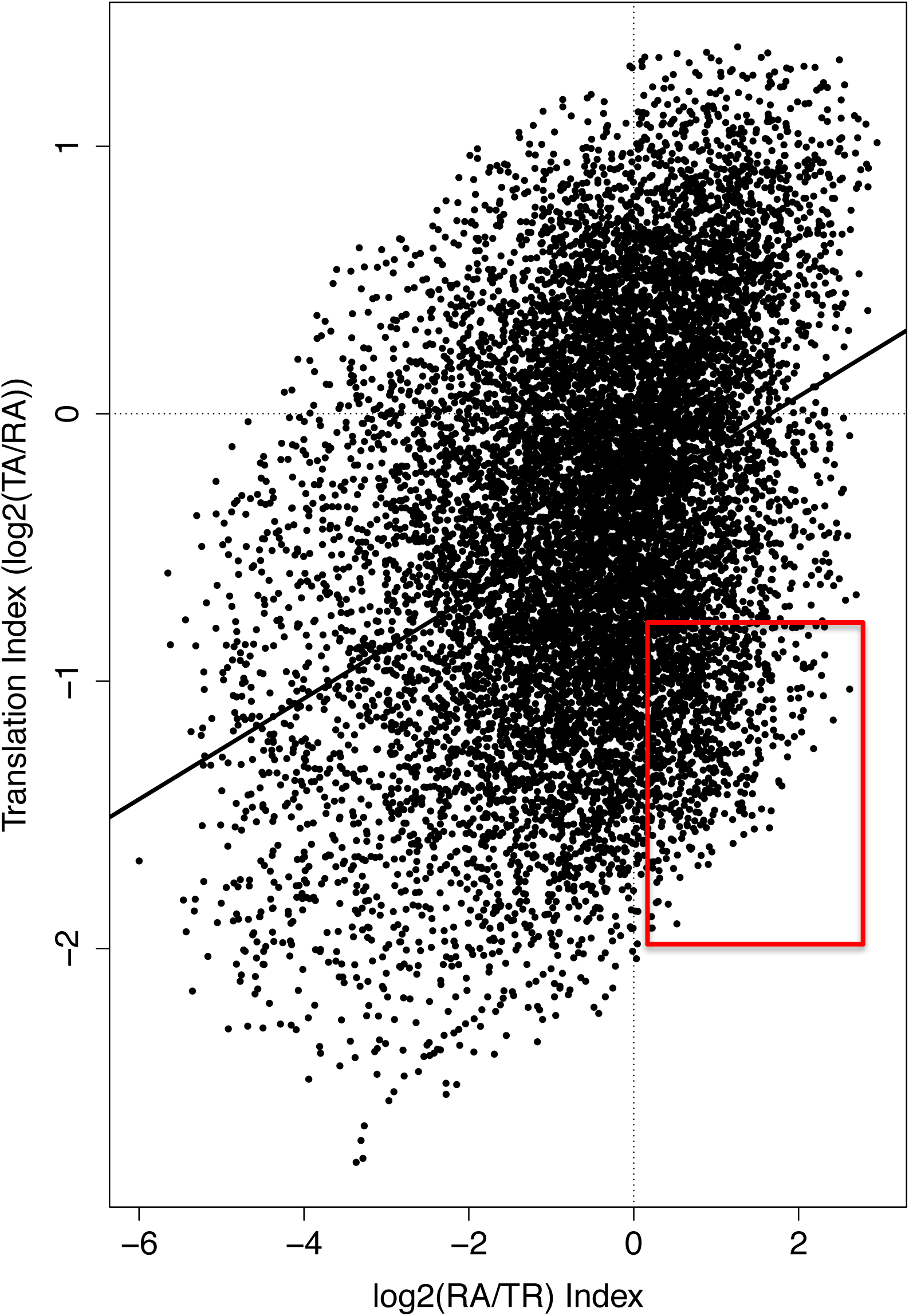
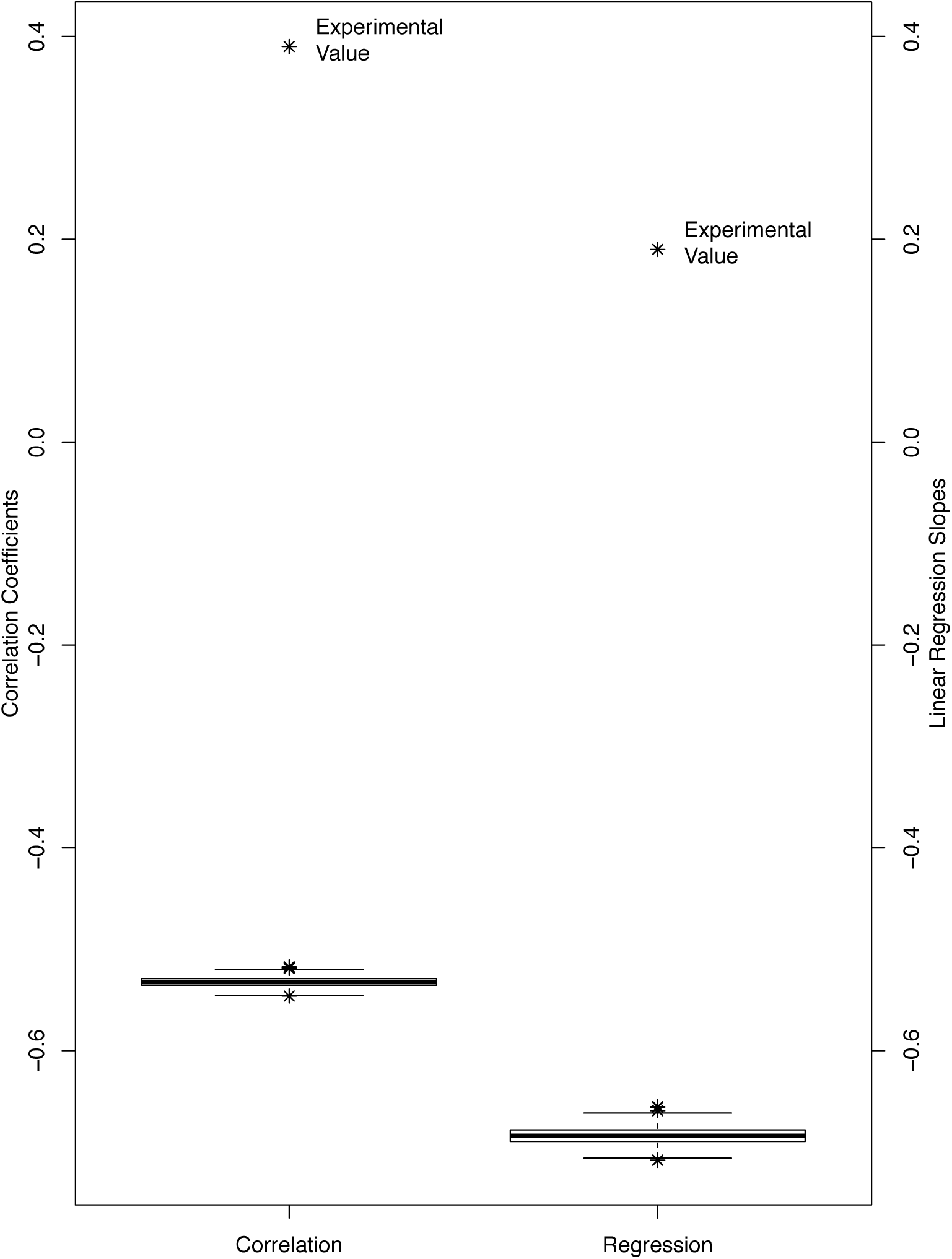
Overall positive correlation between the stability index (log_2_(RA/TR)) and the translation index (log_2_(TA/RA)). A. Scatter plot of the stability index versus the translation index. The correlation coefficient and the linear regression line between the two indices are also shown. The red rectangle identifies mRNAs with high stability but low translation activity to be further analyzed later (see text and Figure 7). B. Box plot of the correlation coefficient and the slope of linear regression lines between the two indexes upon randomization of the experimental dataset. Values from 1000 randomization were used to generate the boxplot. Experimental values are also shown and denoted.

To examine the significance of this observation, we randomly permutated the TR and TA parameters simultaneously to generate a statistical background model. As expected, this led to negative correlation coefficients, and negative slopes of the linear regression line, between the two indexes. We performed the randomization for 1000 times. This generated 1000 correlation coefficients and 1000 slopes of the corresponding linear regression lines, the boxplots of both of which were shown in Figure 4B. Out of the 1000 randomization, not a single positive correlation was observed – both values were always negative. Figure 4B also shows the experimentally determined positive values of the correlation coefficient and the linear regression line slope, demonstrating a sharp contrast with the respective randomly generated values. This contrast illustrates the magnitude of the effect of the stabilization-by-translation regulation mechanism on the relationship between the two indices. Thus, consistent with our hypothesis, the stabilization-by-translation regulatory mechanism renders the relationship into an overall positive one. Moreover, it should be pointed out that the analysis likely under-estimates the significance of the stabilization-by-translation regulatory mechanism, due to the lack of RNA processing information in the TR-RA index.

To examine a potential causal relationship from this stabilization-by-translation regulatory mechanism to the enhancement of gene expression selectivity from transcription to translation, another statistical experiment was performed. We standardized the three genome profiles. This statistical procedure – mean subtraction followed by division by the standard deviation – is routinely used to eliminate statistical differences between distributions. The transformation should, in this case, reduce the numerical differences among the three parameters for many genes shown in figures 2A and 2B; a gene with higher TA and lower TR values than its RA value, for instance, should have similar values for all three parameters in the transformed dataset. That is, the statistical differences among the three genomic profiles shown in figure 3 and table S1 were eliminated by the standardization procedure. If the stabilization-by-translation regulatory mechanism, that is, selective degradation of translationally inactive mRNA, is a major cause of the discrepancy among the three parameters, the positive correlation between the translation and the stability indices must also be significantly reduced in the transformed data. This is, indeed, the case. The correlation coefficient was reduced from 0.39 to 0.12, supporting selective degradation of translationally inactive mRNA as a major cause for the enhancement of gene expression selectivity from transcription to translation. Additionally, many functional genomic normalization procedures rely on standardization of the datasets; and many others, such as the rank and quantile based methods, have similar effects. Our results raised a technical issue that they are not good choices for multi-parameter gene expression study, as valuable information will be lost.

Summarily, all three analysis (correlation, permutation and standardization) support mRNA-stabilization-by-translation as a major contributor to enhancement of gene expression selectivity from transcription to translation, an important functional aspect of transcriptome regulation. This is, to our knowledge, the first quantitative and genome-wide examination of the contribution of this regulatory mechanism.

### Pathway/function specific pattern of correlation between the stability and the translation indices

The correlation between the two indices, on the other hand, is not nearly unequivocal; too many genes deviate significantly from the overall trend – the linear regression line (Fig. 4A). We asked whether this is due to function specific patterns of gene expression regulation, as genes involved in the same biological process have been shown to share a similar pattern in other datasets. To answer this question, we performed two systematic analysis. First, we calculated the distances between the coordinates of each gene pair in Figure 4A. We then created the histograms of the pairwise distances between gene pairs associated with similar sets of gene ontology (GO) terms (see Materials and Methods for detail), and also a histogram for the distances between gene pairs with no significant GO similarity. The distance between genes associated with similar GO terms tend to be smaller than those between genes with no significant similarity in their GO association (Figure 5A). The trend is correlated with the GO similarity score; the higher the score, the more the histogram shifts toward short distance range. Second, we performed this comparison of distances between gene pairs whose proteins interact with each other versus gene pairs whose proteins have not been found to interact with each other. This was done with protein-protein interaction data, which was, as we have previously done [43], downloaded from the IntAct database [44, 45]. As shown in Figure 5B, the interacting pairs exhibit shorter distance than non-interacting pairs. And the trend is correlated with the confidence score assigned to the protein pairs in the IntAct database. Since the protein-protein interaction datasets are generally considered noisy, the confidence score quantify the reliability of the interaction. As shown in Figure 5B, the more reliable the interaction, the more the histogram shifts toward short distance range.

**Figure 5:**
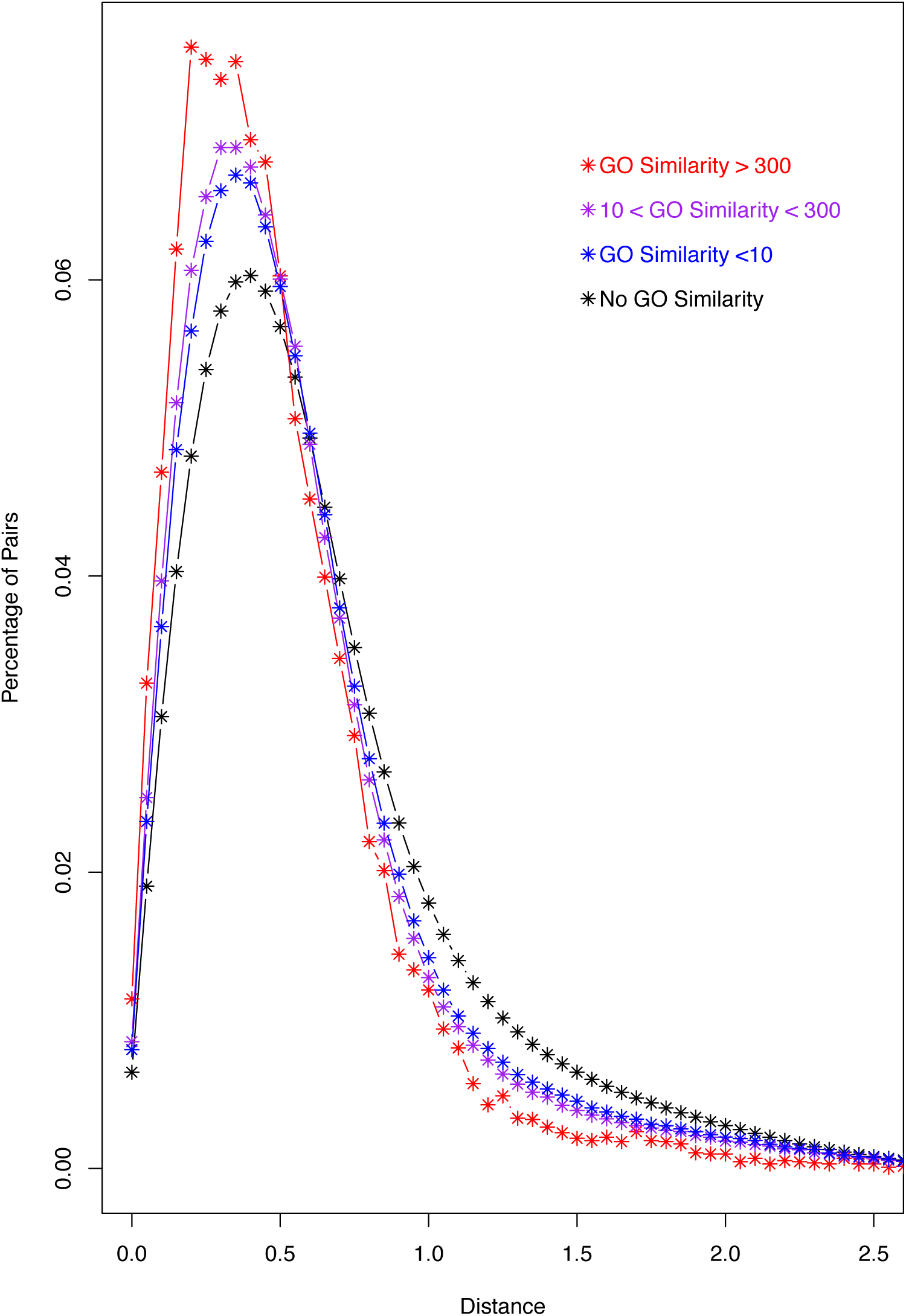
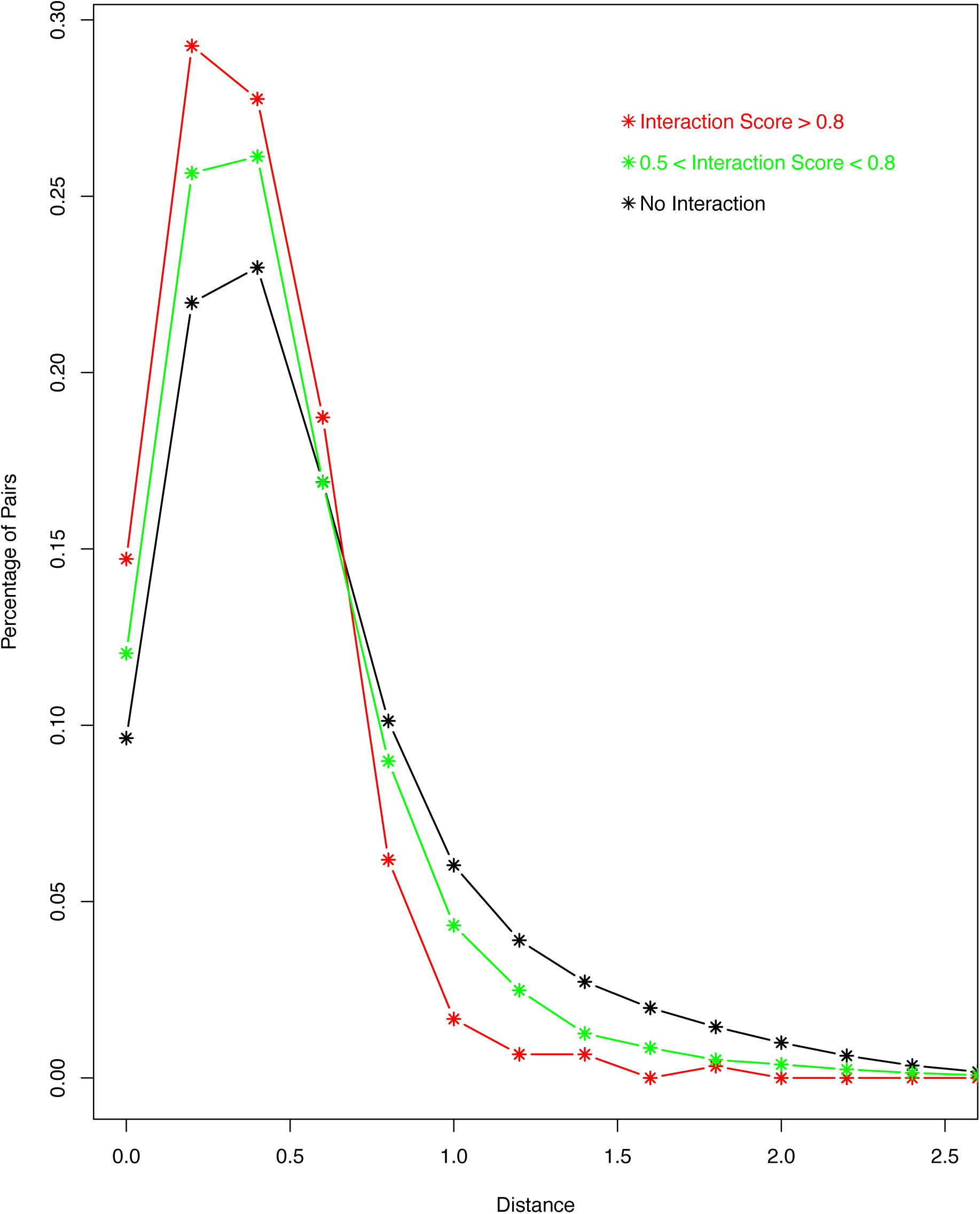
Pathway/function specific pattern of the correlation between the stability and the translation indexes. A. Histograms of the distances between their mRNAs’ coordinates in Figure 3A for gene pairs with different levels of GO similarity. B. Histograms of the distances for gene pairs whose proteins were shown to mutually interact with different levels of interaction confidence.

This function specific pattern is illustrated by the distinct patterns of two exemplary functional groups of genes – the genes for the proteasome subunit (PMS) proteins (PMSA1 to 7 and PMSB1 to 7) and the like-Sm (LSM) genes (LSM1 to 8) (Figure S1). The PMS genes code for proteins that constitute the proteasome 20S core structure [46]. Their mRNAs share a pattern of high levels of both indices. The LSM genes code for subunits of two single-stranded-RNA-binding hetero-heptameric ring structures – one cytoplasmic and the other nucleus [47]. Subunits LSM1 to 7 form the heptamer that is part of the P-body and functions during mRNA degradation in the cytoplasm. Consistently, LSM1-7 mRNAs share a common pattern. However, the pattern is strikingly different from the pattern shared by the PMS mRNAs. While the LSM1-7 mRNAs exhibit relatively high TR-RA index values, unlike PMS mRNA, they exhibit largely lower than average translation activity. The LSM8 subunit interacts with, and nucleus-retains, LSM2 to 7 subunits to form the nucleus heptameric ring structure [47]. That is, it replaces the LSM1 subunit to form the nucleus heptameric structure. This heptameric structure binds to the U6 snRNA and U8 small nucleolar RNA (snoRNA), and thus functions during general RNA maturation in the nucleus. Consistent with this unique LSM8 function, the LSM8 mRNA does not follow the pattern shared by LSM1 to 7 mRNAs, in that it is relatively unstable (Figure S1).

Thus, we observed a function/pathway specific pattern of variation in how much individual mRNA species is regulated by the stabilization-by-translation regulatory mechanism, *i.e.*, the level of correlation between the two indexes; some mRNAs defy this regulation completely in that they stay out of polysome but retain high RA-TR index values and, likely, high stability. Next, we tried to gain mechanistic insight into this variation, and turned our attention to other post-transcription regulatory mechanisms and the untranslated region (UTR) of mRNAs.

### mRNA UTR proportion is a major determinant of whether a mRNA obeys or defies the stabilization-by-translation regulatory mechanism

Besides the stabilization-by-translation regulation, many other mechanisms exist in multi-cellular eukaryotic species, but have not been accounted for in our analysis. For instance, the miRNAs/siRNAs target and regulate a large portion of the transcriptome. Essentially all regulatory signals for such regulation are embedded in mRNA UTR sequences. Consistently, UTRs are abundant in multi-cellular transcriptomes. This is especially true in human. As shown in Figure 6A, on average, the ORF occupies only ∼50% of a human mRNA; the other half is devoted to the UTRs. In many mRNAs, the UTR occupies more than 90% of the total length; for instance, the mRNAs of the all-important CREB1 (cyclic AMP-responsive element-binding protein 1) gene. Since the regulatory signals for mRNA post-transcription regulation are mostly embedded in the UTR sequences, the proportion of an mRNA that is occupied by the UTRs should serve as a good measure of the degree to which the mRNA is controlled by these regulatory mechanisms. Thus, we hypothesized that this proportion should be a major explanatory factor for the high level of variation in the correlation between the two indexes. Indeed, our results support this hypothesis. First, the correlation coefficient between the two indices is optimal at ∼20% UTR, but steadily decreases as this proportion further increases (Fig. 6B); and so is the slope of the linear regression line between the two indices (Fig. 6C). Second, the mRNAs that defy the stabilization-by-translation regulatory mechanism, those that show low translation activity but relatively high RA-TR index values as identified by the red rectangle in Figure 4A, display higher proportion of UTRs. The histogram of their UTR proportions, when compared with that of the whole human transcriptome, shifts clearly toward high proportion ranges (Fig. 7).

**Figure 6:**
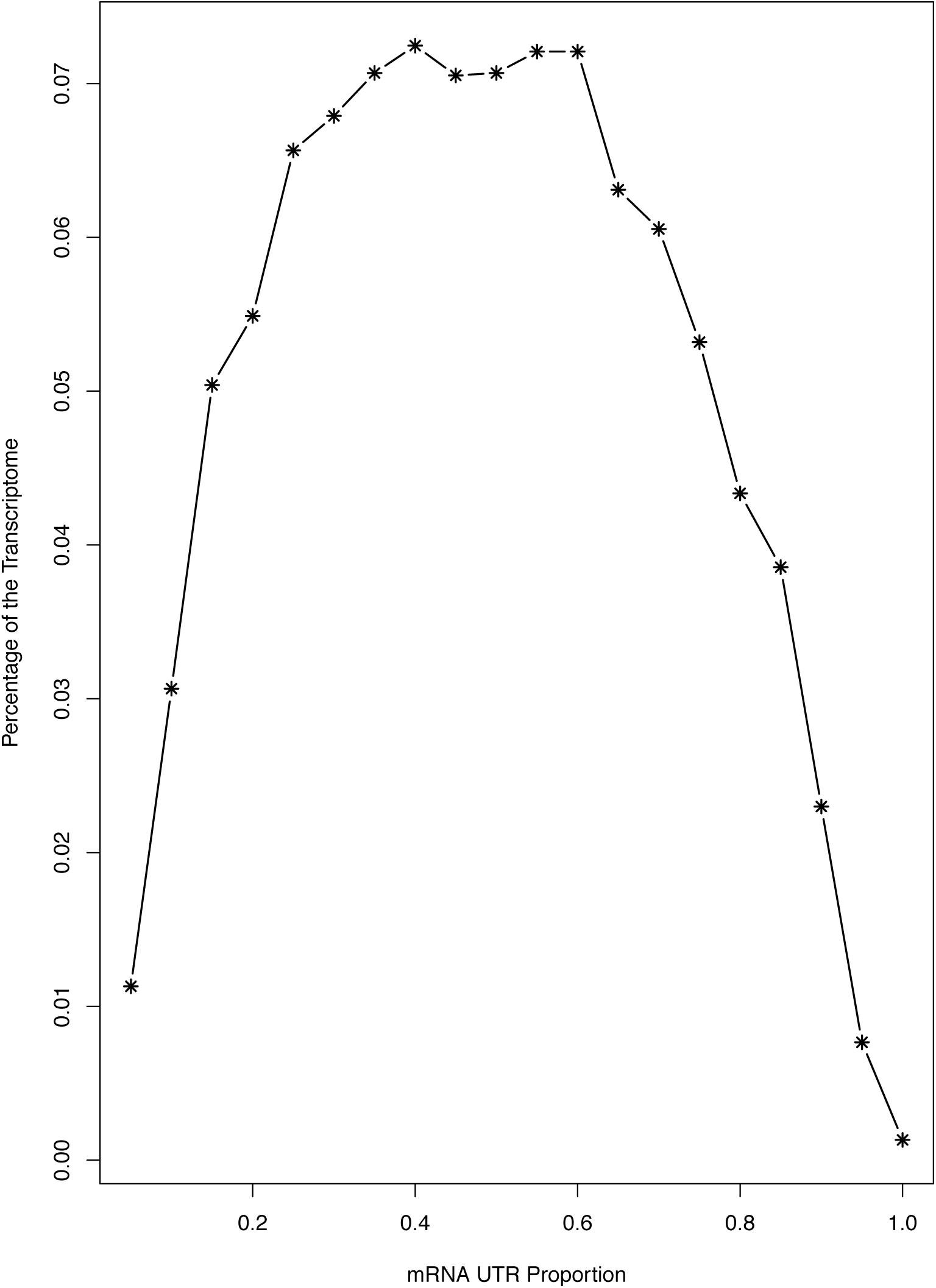
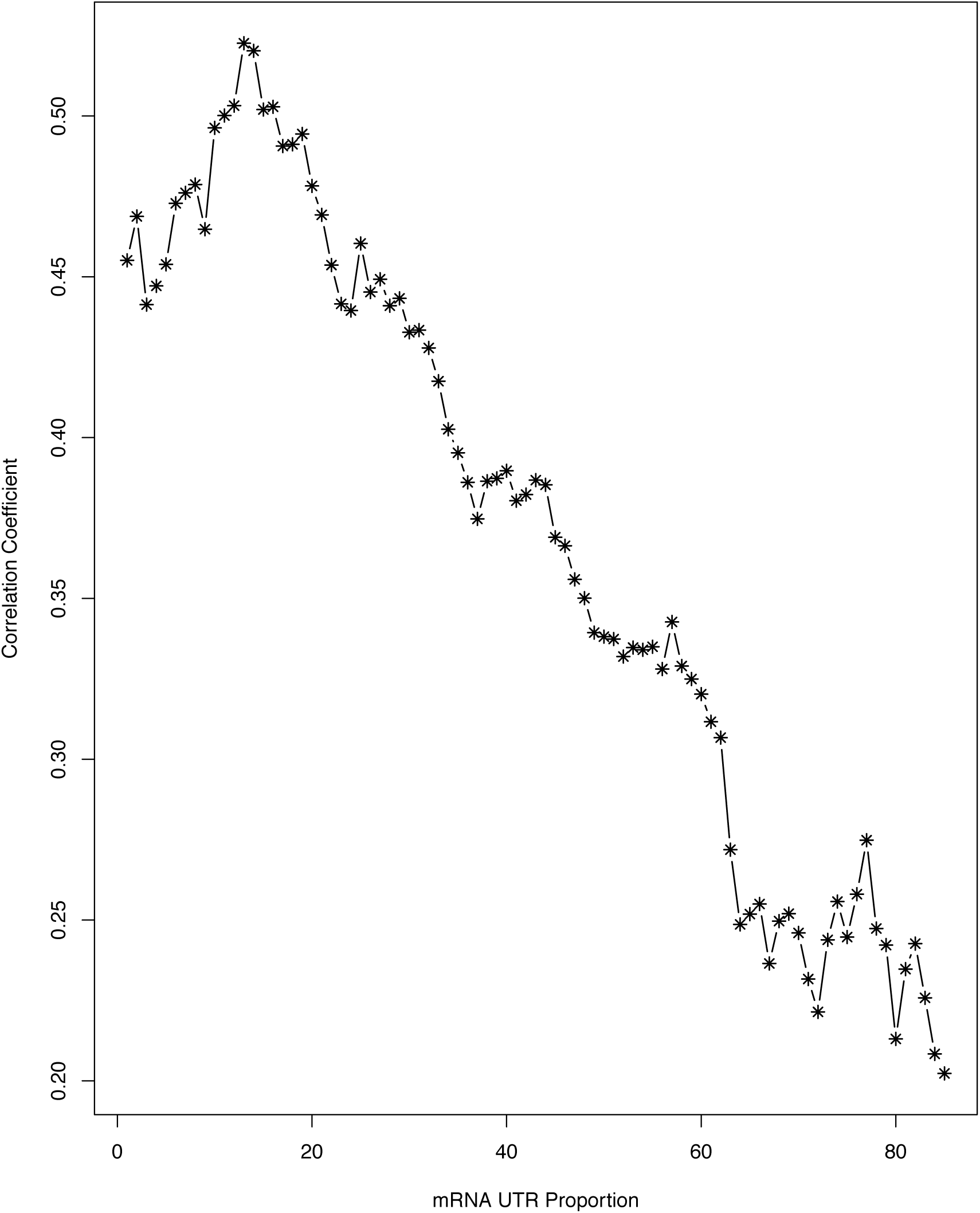
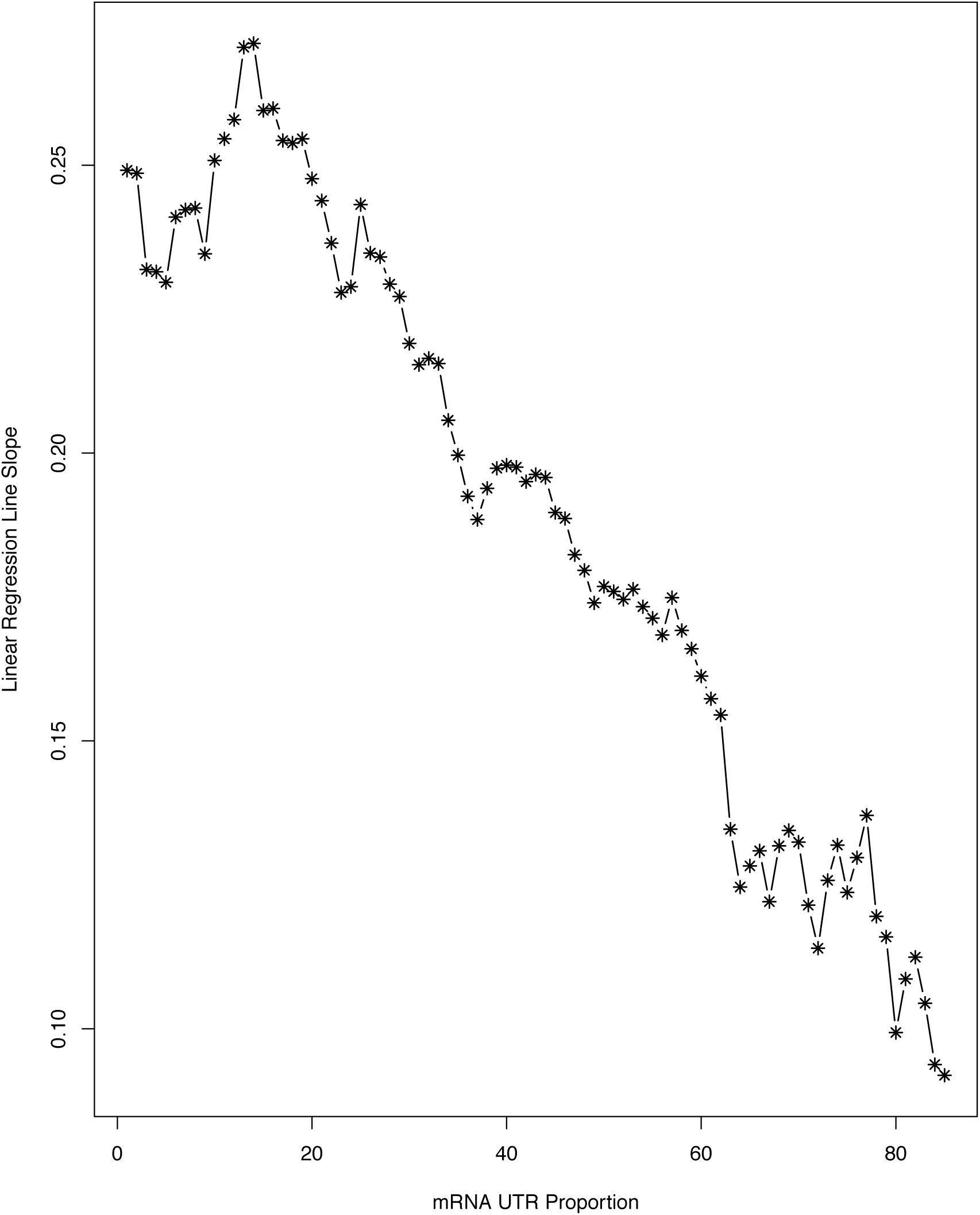
The proportion of a mRNA that is occupied by the UTRs is a determinant of the level of the correlation between the stability and the translation indices. A: Histogram of the UTR proportions of human mRNAs. B and C: The correlation coefficient (B) and the slope of the linear regression line (C) between the stability and the translation indices decrease as the mRNA UTR proportion increases.

**Figure 7:**
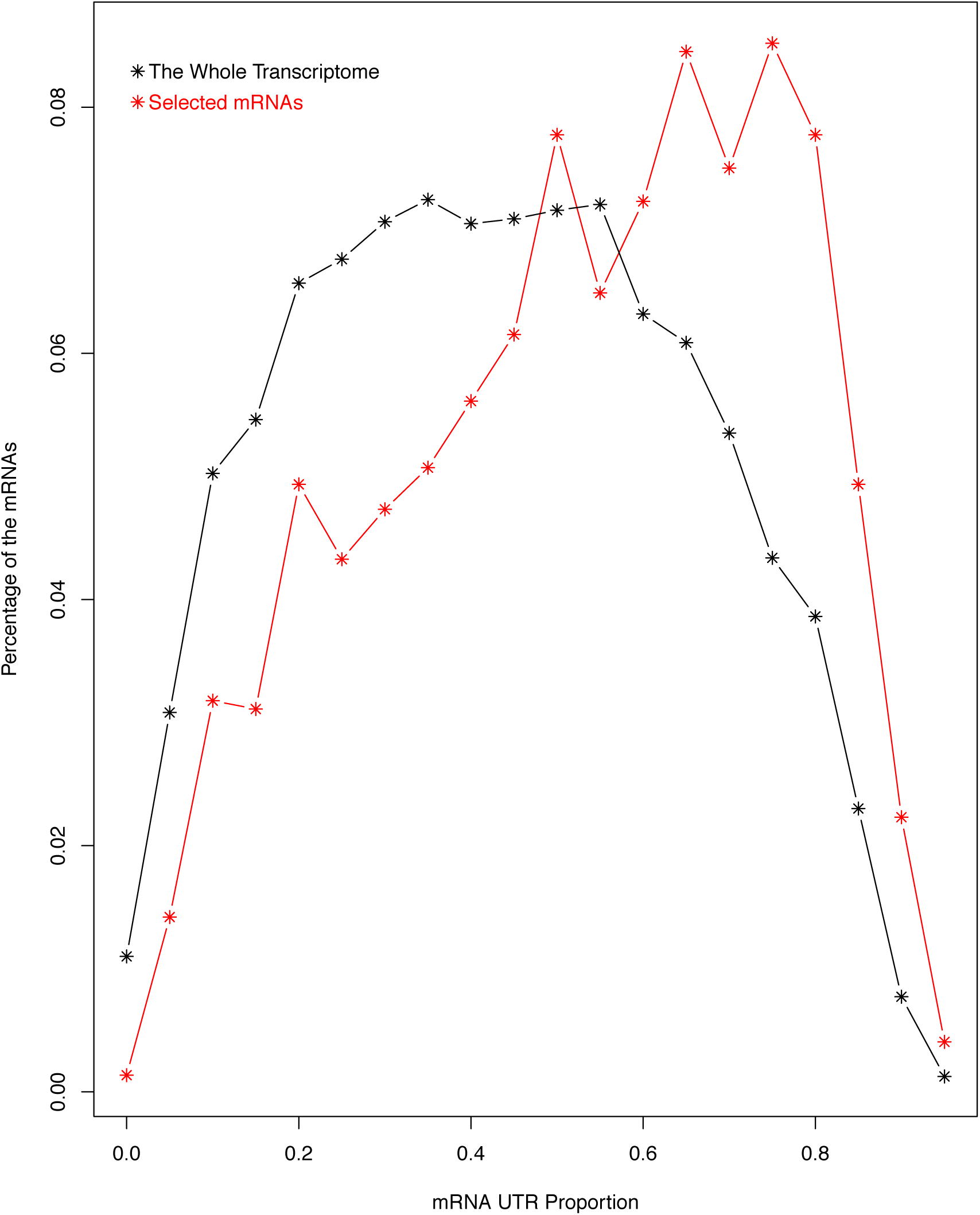
Histogram of the UTR proportions of the mRNAs that have relative high stability but lower-than-average translation activity in comparison with that of the whole human transcriptome. The mRNAs with high stability but low translation activity were selected with the red rectangle in Figure 4A.

To exemplify this observation, we once again used the PMS and the LSM functional groups of genes shown in figure S1 (Table S2). The mRNAs of the PMS group of genes have high levels of both RA-TR index and translation activity, and thus exemplify mRNAs controlled by the stabilization-by-translation mechanism. Consistently, as shown in table 2, they have less-than-average UTR proportions (an average of 35.3% and a median of 26.4%). The mRNAs for the LSM genes, on the other hand, have low levels of translation activity but relatively high RA-TR index values; they thus exemplify mRNAs defying the stabilization-by-translation regulatory mechanism. Not surprisingly, they have higher-than-average UTR proportions (a mean of 69.6% and a median of 68.6%). The difference between the UTR proportions of the mRNAs of the two functional groups of genes is statistically significant; it has, according to a t-test, a p-value of 0.0004 (Table S2).

Thus, variation in the correlation between the two indices can be partially explained by post-transcription regulatory mechanisms mediated by mRNA UTRs. To put it another way, multiple regulatory mechanisms control the transcriptome. Multi-parameter approaches, such as ours as described, have the urgently needed power to dissect cellular co-ordination of these mechanisms and functionally characterize UTR sequences.

## Discussion

Regulation of the transcriptome is a major underpinning of cellular operation. It involves, in addition to transcription, many post-transcriptional processes. Multiple parameters, such as TR, RA and mRNA degradation rate, are relevant to this multi-faceted process. Many multi-parameter studies have been reported and revealed significant discrepancies among the parameters, such as those between TR and RA and those between RA and protein abundance, prompting an appreciation for the complexity of transcriptome regulation.

We have previously participated in the study of the complexity of transcriptome regulation, with a desire for a mechanistic understanding of the discrepancies among TR, RA and protein abundance as well as potential operational advantages the cells gain from them. In this study, we took advantages of the power and versatility of NGS analysis through its coupling to traditional experimental protocols. We simultaneously measured three transcriptome regulation parameters: TR, RA and TA. Based on the close correlation between TR and mRNA production rate, we also indirectly estimate mRNA stability (or degradation rate) by the log ratio of RA and TR (log_2_(RA/TR)). Given the importance of mRNA UTRs in post-transcriptional regulation, the data was analyzed in conjunction with individual mRNAs’ proportions that are UTRs. To put it another way, we broke open the “blackbox” of transcriptome regulation and peeked inside for mechanistic insight into how the cells co-ordinate multiple factors that regulate the transcriptome. In the present paper, we publish, to our knowledge, the first genome-wide dataset that enables integrative analysis of TR, RA, TA and, to some degree, mRNA stability.

It is well known that actively translating mRNA are likely protected from degradation, and thus stabilized. In bacteria, this is considered the primary mechanism for mRNA stability regulation. In eukaryotes, more post-transcriptional regulatory mechanisms, such as micro-RNA control, evolutionarily emerged, giving rise to a more complicated scheme of mRNA stability regulation. However, it is certain that the stabilization-by-translation mechanism still play prominent roles in eukaryotic transcriptome regulation. We provide a quantitative analysis of the impact of translation activity on transcriptome regulation, by showing a moderate but significant positive correlation between the mRNA translation index and the RA-TR index.

This correlation has some explanatory power over the discrepancy between TR and RA. High translation activity protects a mRNA species from degradation, while other less translated mRNAs are being actively degraded and removed out of the transcriptome. This leads to enrichment of the mRNA species, resulting in higher steady-state abundance level than that implied by its production rate and, by extension, TR. Conversely, low translation activity makes a mRNA species more susceptible to the degradation process, leading to situations where the steady-state abundance level is lower than that implied by the production rate. The operational advantages the cells gained by implementing this regulatory scheme remain to be elucidated.

Our multi-parameter approach represents a feasible option to enable the much-needed systematic analysis of mRNA UTRs. The UTRs are much more abundant in the human transcriptome than in any other transcriptome. Their functions in post-transcriptional regulation are well documented. Essentially, all signals for post-transcriptional regulation reside in the UTRs; for instance, microRNA and siRNA target sites, ARE, IRE-IRP etc. But systematic study of mRNA UTRs has been lacking, and our knowledge about their functions remains fragmentary at best. This is perhaps due to a lack of relevant genomic experimental approaches and datasets. mRNA abundance measurement alone is ill-suited for the study of post-transcription regulatory mechanisms and functional analysis of mRNA UTRs. Additionally, though microRNAs/siRNAs target mRNA UTRs and are major regulators of both translation and mRNA degradation, to our knowledge, microRNA/siRNA study has not been integrated with simultaneous genome-wide measurement of translation activity and mRNA stability. Thus, our integrative multi-parameter analysis represents a novel functional genomic approach to mRNA UTR analysis. It is able to reveal that the UTRs potentially play an important role in maintaining the stability of translationally inactive mRNA species, thus conferring to human cells the capacity for a post-transcriptional regulatory mechanism that is absent in prokaryotic species and mostly in uni-cellular species such as the yeast *S. cerevisiae*. It should be noted that our results represent only a single time-point snap-shot of actively growing human cells. More power of this analysis approach, we believe, is yet to be relished in analyzing dynamic changes of the three parameters during physiological processes, and when it is coupled to more accurate mRNA production rate measurement techniques such as nascent-RNA-metabolic-labeling based approaches.

Additionally, computational analysis of mRNA UTRs for key regulatory signals embedded in the UTR sequences remains technically challenging. This is due to low signal-to-noise ratio and the lack of a general guiding principle. For instance, a typical mammalian microRNA target site is no more than 8 nucleotide long. Our approach provides a way to classify the mRNAs based on their patterns in the generated datasets, i.e., their behavior in the multi-faceted transcriptome regulation process. Key regulatory signals shall be shared by the UTRs of similarly classified mRNAs, and thus can be computationally extracted from them – a much easier approach than *de novo* computational analysis of mRNA UTR sequences. That is, datasets generated through this approach should provide a functional context for enhancing the signal-to-noise ratio in computational analysis of mRNA UTR sequences.

We also quantitatively describe the trend of sequentially higher levels of selectivity as the genetic information flow from the genome to the proteome in the gene expression process. In other words, the gene expression machinery focuses its resources on less and less genes, so that only mission critical proteins are expressed in the proteome. The multi-stepped gene expression process can be considered as, to some degree, a selective amplification process. Transcription selectively amplifies the genomic sequences into multiple copies of mRNA sequences. Translation, in turn, selectively amplifies individual mRNA molecules into multiple copies of protein sequences. The selectivity of this process is further enhanced by selective mRNA degradation. Even though obvious from the results in previous publications, this trend of sequentially higher levels of selectivity in the gene expression process has not received much attention, and was never explicitly stated in these reports. In this study, we quantitatively described this trend by comparing the dispersions of the genomic profiles of the three gene expression parameters. Our results also suggest that mRNA degradation plays perhaps the biggest role in this trend, as the jump in selectivity from transcription rate to mRNA abundance is much bigger than the increase from mRNA abundance to translation activity. That is, selective degradation of those mRNAs, which are not protected from degradation by active translation or other processes mediated by their UTRs, play an important role in shaping up the transcriptome and priming it for efficient production of mission-critical proteins.

## Conclusions

In summary, we present a quantitative delineation of cellular coordination of transcription, mRNA abundance, mRNA translation activity, to some degree mRNA stability as well as mechanistic involvement of mRNA UTRs in the coordination process. As a consequence of the coordination activity, the cells exhibit sequentially higher level of gene expression selectivity from transcription to mRNA abundance, and then to translation activity. The results contribute to our understanding of the complexity of the multi-stepped gene expression process, through which the cells “read” the genomic “book” of seemingly simplistic string of nucleotides and “translate” information embedded in the sequences into cellular operations, that is, dynamic control of the biochemical flow through biochemical reactions, pathways and networks [1-4].

## Methods

### Tissue Culture and mRNA Isolation for RNA-seq Analysis

The human HCT116 cells were cultured in a serum-free medium (McCoy’s 5A (Sigma) with pyruvate, vitamins, amino acids and antibiotics) supplemented with 10 ng/ml epidermal growth factor, 20 µg/ml insulin and 4 µg/ml transferrin. Cells were maintained at 37 °C in a humidified incubator with 5% CO_2_.

To extract mRNA for RNA-seq analysis, RNeasy kit (Qiagen) was used to extract total RNA from the HCT116 cells according to manufacture’s specification. GeneRead Pure mRNA Kit (Qiagen) was then used to isolate mRNA from the total RNA for Illumina NGS sequencing according to manufacture’s specification.

### GRO-seq Analysis

Global run-on was done as previously described [26, 48, 49]. Briefly, two 100cm plates of HCT116 cells were washed 3 times with cold PBS buffer. Cells were then swelled in swelling buffer (10mM Tris-pH7.5, 2mM MgCl2, 3mM CaCl2) for 5min on ice. Harvested cells were re-suspended in 1ml of the lysis buffer (swelling buffer with 0.5% IGEPAL and 10% glycerol) with gentle vortex and brought to 10ml with the same buffer for nuclei extraction. Nuclei were washed with 10ml of lysis buffer and re-suspended in 1ml of freezing buffer (50mM Tris-pH8.3, 40% glycerol, 5mM MgCl2, 0.1mM EDTA), pelleted down again, and finally re-suspended in 100µl of freezing buffer.

For the nuclear run-on step, re-suspended nuclei were mixed with an equal volume of reaction buffer (10mM Tris-pH 8.0, 5mM MgCl2, 1mM DTT, 300mM KCl, 20 units of SUPERase-In, 1% Sarkosyl, 500µM ATP, GTP, and Br-UTP, 2µM CTP) and incubated for 5 min at 30°C. Nuclei RNA were extracted with TRIzol LS reagent (Invitrogen) following manufacturer’s instructions, and was resuspended in 20µl of DEPC-water. RNA was then purified through a p-30 RNAse-free spin column (BioRad), according to the manufacturer’s instructions and treated with 6.7µl of DNase buffer and 10µl of RQ1 RNase-free DNase (Promega), purified again through a p-30 column. A volume of 8.5µl 10×antarctic phosphatase buffer, 1µl of SUPERase-In, and 5µl of antarctic phosphatase was added to the run-on RNA and treated for 1hr at 37°C. Before proceeding to immuno-purification, RNA was heated to 65°C for 5min and kept on ice.

Anti-BrdU argarose beads (Santa Cruz Biotech) were blocked in blocking buffer (0.5×SSPE, 1mM EDTA, 0.05% Tween-20, 0.1% PVP, and 1mg/ml BSA) for 1 hr at 4°C. Heated run-on RNA (∼85µl) was added to 60µl beads in 500µl binding buffer (0.5×SSPE, 1mM EDTA, 0.05% Tween-20) and allowed to bind for 1hr at 4°C with rotation. After binding, beads were washed once in low salt buffer (0.2×SSPE, 1mM EDTA, 0.05% Tween-20), twice in high salt buffer (0.5% SSPE, 1mM EDTA, 0.05% Tween-20, 150mM NaCl), and twice in TET buffer (TE pH7.4, 0.05% Tween-20). BrdU-incorporated RNA was eluted with 4×125µl elution buffer (20mM DTT, 300mM NaCl, 5mM Tris-pH 7.5, 1mM EDTA, and 0.1% SDS). RNA was then extracted with acidic phenol/chloroform once, chloroform once and precipitated with ethanol overnight. The precipitated RNA was re-suspended in 50µl reaction (45µl of DEPC water, 5.2µl of T4 PNK buffer, 1µl of SUPERase_In and 1µl of T4 PNK (NEB)) and incubated at 37°C for 1 hr. The RNA was extracted and precipitated again as above before being processed for Illumina NGS sequencing.

### Polysome Isolation and mRNA extraction

Polysome was isolated as previously described [50, 51]. Briefly, the HCT116 cells were incubated with 100µg/ml cycloheximide for 15 minutes, washed three times with PBS, scraped off into PBS, and then pelleted by micro-centrifugation. Cell pellet was homogenized in a hypertonic re-suspension buffer (10 mM Tris (pH 7.5), 250 mM KCl, 2 mM MgCl_2_ and 0.5% Triton X100) with RNAsin RNAse inhibitor and a protease cocktail. Homogenates were centrifuged for 10 min at 12,000 g to pellet the nuclei. The post-nuclear supernatants were laid on top of a 10-50% (w/v) sucrose gradient, followed by centrifugation for 90 min at 200,000 g. The polysomal fractions were identified by OD_254_ and collected. RNeasy kit (Qiagen) was used to extract RNA from the polysome fractions according to manufacture’s specification. GeneRead Pure mRNA Kit (Qiagen) was then used to isolate mRNA for Illumina NGS sequencing from the RNA according to manufacture’s specification.

### Illumina NGS Sequencing

Sequencing libraries was generated with the Illumina TruSeq RNA Sample Preparation Kit. Briefly, RNA molecules were fragmented into small pieces using divalent cations under elevated temperature. The cleaved RNA fragments are copied into first strand cDNA synthesis using reverse transcriptase and random primers. This was followed by second strand cDNA synthesis using DNA Polymerase I and RNase H. These cDNA fragments were end-repaired using T4 DNA polymerase, Klenow polymerase and T4 polynucleotide kinase. The resulting blunt-ended fragments were A-tailed using a 3′–5′ exonuclease-deficient Klenow fragment and ligated to Illumina adaptor oligonucleotides in a ‘TA’ ligation. The ligation mixture was further size-selected by AMPure beads and enriched by PCR amplification following Illumina TruSeq DNA Sample Preparation protocol. The resulting library is attached and amplified on a flow-cell by cBot Cluster Generation System.

The sequencing was done with an Illumina HiSeq 2000 sequencer. Multiplexing was used to pool 4 samples into one sequencing lane. After each sequencing run, the raw reads were pro-processed to filter out low quality reads and to remove the multiplexing barcode sequences.

### NGS Data Analysis

The sequencing reads were mapped to the UCSC hg19 human genome sequences with the TopHat software, using the default input parameter values. For each sample, at least 80% of the reads were successfully mapped. For the sake of consistence across the three transcriptome regulation parameters, we counted the reads for each gene for the exon regions only. The counting was performed with the HTSeq-count software, and the counts were then transformed into Reads Per Kilo-base Per Million Mapped Reads (RPKM) values. 12921 genes have a minimal RPKM value of 1 for at least one of the three parameters, and were considered expressed in the HCT116 cells. Linear regression of log transformed data was used to examine consistence between biological replicate samples.

### Statistical Analysis

The R open source statistical software (version 3.3.1) installed on a Mac Pro desktop computer was used for statistical analysis. Outlier identification, student t-test, standard deviation calculation, correlation coefficient calculation, linear regression and other statistical procedure are all done with this R software.

The procedure for comparing the linear regression slopes/coefficients shown in figure 2B is described as follows. We first applied the following linear regression models to the data:

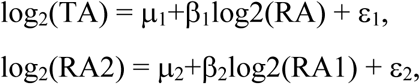

and ε_1_ and ε_2_ follow normal distribution.

It is estimated that

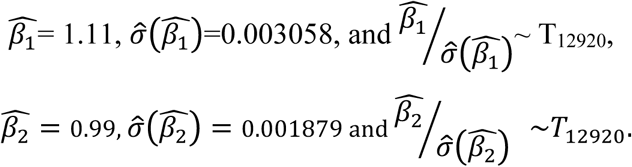

Therefore, the 97.5% confidence interval for *β*_1_ is

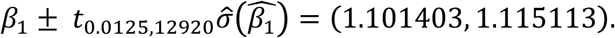

The 97.5% confidence interval for *β*_2_ is

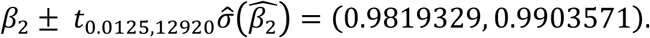

These two confidence intervals do not overlap, implying that, at significant level 0.05, the two regression coefficients are different.

In addition, because T distribution with degrees of freedom of 12920 is very close to standard normal distribution, the t-score, calculated as below,

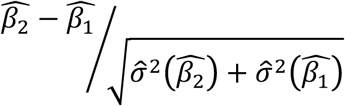

approximately follow standard normal distribution. This allows p-value calculation. The p-value is essentially 0 (smaller than 1E-200).

### Gene Ontology (GO) Similarity Analysis

Pairwise GO similarity score between human genes was computed as previously described [52-54]. Briefly, for each gene, we first generated GO fingerprint – a set of ontology terms enriched in the PubMed abstracts linked to the gene, along with the adjusted p-value reflecting the degree of enrichment of each term. The GO similarity score quantifies similarity between the GO fingerprints of corresponding gene pair. For detail about GO fingerprint generation and similarity calculation, please see description in previous publications [53].

## Declaration

### Ethics approval

Not applicable.

### Consent for publication

Not applicable.

### Availability of data and materials

The dataset is being deposited into the NCBI GEO database. Accession number will be provided once it is available.

### Authors’ contributions

WJ performed molecular and cellular experimental portion of the project. ZG, NL, WJZ and FZ performed computational and statistical analysis. DF performed polysome isolation. WJ and DG designed the project and drafted the manuscript. All authors participated in critical revision of the manuscript and have given final approval for publication.

## Acknowledgements

Not applicable.

## References

1. Searls, D.B., Linguistic approaches to biological sequences. Comput Appl Biosci, 1997. 13 (9283748): p. 333–344.

2. Searls, D.B., Reading the book of life. Bioinformatics, 2001. 17(7): p. 579–80.

3. Searls, D.B., The language of genes. Nature, 2002. 420(12432405): p. 211–217.

4. Wang, D.G., “Molecular gene”: Interpretation in the right context. Biology & Philosophy, 2005. 20(2-3): p. 453–464.

5. Jovanovic, M., et al., Immunogenetics. Dynamic profiling of the protein life cycle in response to pathogens. Science, 2015. 347(6226): p. 1259038.

6. Rabani, M., et al., High-resolution sequencing and modeling identifies distinct dynamic RNA regulatory strategies. Cell, 2014. 159(7): p. 1698–710.

7. Li, J.J., P.J. Bickel, and M.D. Biggin, System wide analyses have underestimated protein abundances and the importance of transcription in mammals. PeerJ, 2014. 2: p. e270.

8. Liu, Y. and R. Aebersold, The interdependence of transcript and protein abundance: new data--new complexities. Mol Syst Biol, 2016. 12(1): p. 856.

9. Schwanhausser, B., et al., Global quantification of mammalian gene expression control. Nature, 2011. 473(7347): p. 337–42.

10. McManus, J., Z. Cheng, and C. Vogel, Next-generation analysis of gene expression regulation-comparing the roles of synthesis and degradation. Mol Biosyst, 2015. 11(10): p. 2680–9.

11. Vogel, C. and E.M. Marcotte, Insights into the regulation of protein abundance from proteomic and transcriptomic analyses. Nat Rev Genet, 2012. 13(4): p. 227–32.

12. Anderson, L. and J. Seilhamer, A comparison of selected mRNA and protein abundances in human liver. Electrophoresis, 1997. 18(9150937): p. 533–537.

13. Gygi, S.P., et al., Correlation between protein and mRNA abundance in yeast. Mol Cell Biol, 1999. 19(10022859): p. 1720–1730.

14. Ideker, T., et al., Integrated genomic and proteomic analyses of a systematically perturbed metabolic network. Science, 2001. 292(11340206): p. 929–934.

15. Flory, M.R., et al., Quantitative proteomic analysis of the budding yeast cell cycle using acid-cleavable isotope-coded affinity tag reagents. Proteomics, 2006. 6(17133367): p. 6146–6157.

16. Ghaemmaghami, S., et al., Global analysis of protein expression in yeast. Nature, 2003. 425 (6959): p. 737–41.

17. Griffin, T.J., et al., Complementary profiling of gene expression at the transcriptome and proteome levels in Saccharomyces cerevisiae. Molecular & cellular proteomics: MCP, 2002. 1(4): p. 323–33.

18. Le Roch, K.G., et al., Global analysis of transcript and protein levels across the Plasmodium falciparum life cycle. Genome Res, 2004. 14(11): p. 2308–18.

19. Tian, Q., et al., Integrated genomic and proteomic analyses of gene expression in Mammalian cells. Mol Cell Proteomics, 2004. 3(10): p. 960–9.

20. Washburn, M.P., et al., Protein pathway and complex clustering of correlated mRNA and protein expression analyses in Saccharomyces cerevisiae. Proc Natl Acad Sci U S A, 2003. 100(12626741): p. 3107–3112.

21. Cheng, Z., et al., Differential dynamics of the mammalian mRNA and protein expression response to misfolding stress. Mol Syst Biol, 2016. 12(1): p. 855.

22. Garcia-Martinez, J., A. Aranda, and J.E. Perez-Ortin, Genomic run-on evaluates transcription rates for all yeast genes and identifies gene regulatory mechanisms. Mol Cell, 2004. 15(15260981): p. 303–313.

23. Marin-Navarro, J., et al., Global estimation of mRNA stability in yeast. Methods Mol Biol, 2011. 734: p. 3–23.

24. Molina-Navarro, M.M., et al., Comprehensive transcriptional analysis of the oxidative response in yeast. J Biol Chem, 2008. 283(18424442): p. 17908–17918.

25. Romero-Santacreu, L., et al., Specific and global regulation of mRNA stability during osmotic stress in Saccharomyces cerevisiae. RNA, 2009. 15(19369426): p. 1110–1120.

26. Core, L.J., J.J. Waterfall, and J.T. Lis, Nascent RNA sequencing reveals widespread pausing and divergent initiation at human promoters. Science, 2008. 322(5909): p. 1845–8.

27. Hah, N., et al., A rapid, extensive, and transient transcriptional response to estrogen signaling in breast cancer cells. Cell, 2011. 145(4): p. 622–34.

28. Eser, P., et al., Periodic mRNA synthesis and degradation co-operate during cell cycle gene expression. Mol Syst Biol, 2014. 10: p. 717.

29. Rabani, M., et al., Metabolic labeling of RNA uncovers principles of RNA production and degradation dynamics in mammalian cells. Nat Biotechnol, 2011. 29(5): p. 436–42.

30. Dolken, L., et al., High-resolution gene expression profiling for simultaneous kinetic parameter analysis of RNA synthesis and decay. RNA, 2008. 14(9): p. 1959–72.

31. Friedel, C.C., et al., Conserved principles of mammalian transcriptional regulation revealed by RNA half-life. Nucleic Acids Res, 2009. 37(17): p. e115.

32. Brar, G.A., et al., High-resolution view of the yeast meiotic program revealed by ribosome profiling. Science, 2012. 335(6068): p. 552–7.

33. Greenbaum, D., et al., Comparing protein abundance and mRNA expression levels on a genomic scale. Genome Biol, 2003. 4(12952525): p. 117–117.

34. Ingolia, N.T., et al., Genome-wide analysis in vivo of translation with nucleotide resolution using ribosome profiling. Science, 2009. 324(5924): p. 218–23.

35. Coldwell, M.J., N.K. Gray, and M. Brook, Cytoplasmic mRNA: move it, use it or lose it! Biochem Soc Trans, 2010. 38(6): p. 1495–9.

36. Morozov, I.Y., et al., mRNA 3’ tagging is induced by nonsense-mediated decay and promotes ribosome dissociation. Mol Cell Biol, 2012. 32(13): p. 2585–95.

37. Hayles, B., S. Yellaboina, and D. Wang, Comparing transcription rate and mRNA abundance as parameters for biochemical pathway and network analysis. PloS one, 2010. 5(3): p. e9908.

38. Wang, D., Discrepancy between mRNA and protein abundance: insight from information retrieval process in computers. Computational biology and chemistry, 2008. 32(6): p. 462–8.

39. Goodwin, S., J.D. McPherson, and W.R. McCombie, Coming of age: ten years of next-generation sequencing technologies. Nat Rev Genet, 2016. 17(6): p. 333–51.

40. Kim, D., et al., TopHat2: accurate alignment of transcriptomes in the presence of insertions, deletions and gene fusions. Genome Biol, 2013. 14(4): p. R36.

41. Anders, S., P.T. Pyl, and W. Huber, HTSeq--a Python framework to work with high-throughput sequencing data. Bioinformatics, 2015. 31(2): p. 166–9.

42. Pandit, S., D. Wang, and X.D. Fu, Functional integration of transcriptional and RNA processing machineries. Curr Opin Cell Biol, 2008. 20(3): p. 260–5.

43. Guo, Z., et al., Relationship between gene duplicability and diversifiability in the topology of biochemical networks. BMC Genomics, 2014. 15: p. 577.

44. Orchard, S., et al., The MIntAct project--IntAct as a common curation platform for 11 molecular interaction databases. Nucleic Acids Res, 2014. 42(Database issue): p. D358–63.

45. Kerrien, S., et al., The IntAct molecular interaction database in 2012. Nucleic acids research, 2012. 40(Database issue): p. D841–6.

46. Kish-Trier, E. and C.P. Hill, Structural biology of the proteasome. Annu Rev Biophys, 2013. 42: p. 29–49.

47. Khusial, P., R. Plaag, and G.W. Zieve, LSm proteins form heptameric rings that bind to RNA via repeating motifs. Trends Biochem Sci, 2005. 30(9): p. 522–8.

48. Jin, F., et al., A high-resolution map of the three-dimensional chromatin interactome in human cells. Nature, 2013. 503(7475): p. 290–4.

49. Wang, D., et al., Reprogramming transcription by distinct classes of enhancers functionally defined by eRNA. Nature, 2011. 474(7351): p. 390–4.

50. Day, R.T., et al., Acute hyperglycemia rapidly stimulates VEGF mRNA translation in the kidney. Role of angiotensin type 2 receptor (AT2). Cell Signal, 2010. 22(12): p. 1849–57.

51. Feliers, D., et al., Translational regulation of vascular endothelial growth factor expression in renal epithelial cells by angiotensin II. Am J Physiol Renal Physiol, 2005. 288(3): p. F521–9.

52. Qin, T., et al., Finding pathway-modulating genes from a novel Ontology Fingerprint-derived gene network. Nucleic Acids Res, 2014. 42(18): p. e138.

53. Qin, T., et al., Signaling network prediction by the Ontology Fingerprint enhanced Bayesian network. BMC Syst Biol, 2012. 6 Suppl 3: p. S3.

54. Tsoi, L.C., et al., Evaluation of genome-wide association study results through development of ontology fingerprints. Bioinformatics, 2009. 25(10): p. 1314–20.

